# Constructing structural and functional brain networks from regional brain profiles

**DOI:** 10.1101/2025.01.08.632053

**Authors:** Donghui Song

## Abstract

Brain network analysis has become an important approach to understanding brain function. Given the human brain’s complexity and dynamic nature, traditional brain networks based on temporal synchronization may not capture all the nuances of brain activity. The emerging trend is to construct brain networks from regional information, creating pathways that connect regional profiles to network-level insights to better understand brain function. In this study, we constructed structural networks based on regional brain structural information, specifically gray matter volume (GMV), and functional networks based on regional brain activity, measured by brain entropy (BEN). We compared these newly constructed networks with traditional networks: functional connectivity networks (FCN) based on temporal synchronization, structural connectivity networks (SCN) derived from fiber tracking, and MEG-derived functional networks. Our results reveal that GMV network (GMVN) and BEN network (BENN) show correlations with these networks but also exhibit distinct differences. Furthermore, we conducted connectome gradient analyses, uncovering meaningful brain function distribution patterns in both GMVN and BENN. Finally, we used GMVN, BENN, and traditional FCN to predict cognitive and emotional scores. The results showed that BENN, in the resting state, provided the best prediction of both cognitive and emotional scores.

This study systematically evaluates the relationship between brain networks constructed from regional brain information and existing networks, as well as their ability to predict behavioral phenotypes. It demonstrates that networks built from regional information capture aspects of brain activity that traditional networks cannot represent and may even provide superior predictive power for behavioral phenotypes. This opens a new path for understanding brain function from regional to network-level information and lays the foundation for future applications in brain development, individual differences, and clinical research.

## 1. Introduction

The human brain, composed of approximately one hundred billion interconnected neurons, represents an extraordinarily complex system. In recent years, with advancements in network science and neuroscience, network neuroscience has emerged as a cutting-edge approach to understanding brain function. Traditionally, the construction of large-scale brain networks has relied on structural networks derived from fiber tracking using diffusion MRI (dMRI) and functional networks based on temporal synchrony using functional MRI (fMRI). However, these conventional methods may not fully capture the intricacies of complex brain functions. In this context, we propose a novel approach to brain network construction that aims to complement and enhance existing methodologies, providing new insights into the dynamic and multifaceted nature of brain function. This novel brain network construction method is primarily based on the covariation of regional brain profiles. In this study, we focus on two key regional brain profiles: grey matter volume (GMV) for the structural network, and regional brain entropy (rBEN) for the functional network.

GMV was assessed by voxel-based morphometry (VBM), which is one of the most widely used computational approaches for anatomical brain image analysis (Ashburner and Friston 2000, Mechelli, Price et al. 2005). GMV and GMV-derived structural covariance networks (Evans 2013) have been widely employed in brain development and brain disorders (Brickman, Habeck et al. 2007, Garrido, Furl et al. 2009, Groeschel, Vollmer et al. 2010, Montembeault, Joubert et al. 2012, Alexander-Bloch, Giedd et al. 2013, Wise, Radua et al. 2017, Ancelin, Carrière et al. 2019, Kandilarova, Stoyanov et al. 2019, Wang, Yang et al. 2020, Bethlehem, Seidlitz et al. 2022, Christova and Georgopoulos 2023). Our recent research has revealed that structural covariance networks exhibit characteristics similar to those of traditional structural and functional connectivity networks (Song and Wang 2024). Specifically, the similarity within large-scale brain networks is greater than that observed between networks, suggesting that individual-based GMV covariance networks (GMVN) may carry meaningful information.

Brain entropy (BEN) reflects the irregularity, disorder, and complexity of brain activity (Wang, Li et al. 2014). The fMRI-based rBEN has established its distribution in the normal brain (Wang, Li et al. 2014, Song and Wang 2024) and revealed a relationship between cognitive function (Wang 2021, Del Mauro and Wang 2024) and task performance (Lin, Chang et al. 2022, Camargo, Del Mauro et al. 2024, Song and Wang 2024). More importantly, BEN also reflects the effects of neurochemical signals (Song and Wang 2024, Song and Wang 2024, Song and Wang 2024) and neuroplasticity (Chang, Zhang et al. 2018, Song, Chang et al. 2019, Liu, Song et al. 2024, Song, Deng et al. 2024) and has been shown to be associated with various brain disorders (Zhou, Zhuang et al. 2016, Xue, Yu et al. 2019, Liu, Song et al. 2020, Wang and Initiative 2020, Jiang, Cai et al. 2023, Song Donghui 2023, Del Mauro, Sevel et al. 2024, Dong-Hui Song 2024). Recently we also have identified the relationship between regional BEN and brain networks (Song 2024). However, it remains unclear whether rBEN can be used to construct meaningful functional brain networks at an individual level. In previous studies, rBEN networks (rBENN) were commonly constructed based on the co-variation of rBEN across individuals (Wang, Li et al. 2014, Song and Wang 2024), like the construction of structural covariance networks (Evans 2013, Carmon, Heege et al. 2020). The networks, built on the co-variation of rBEN, exhibited a distribution like that observed in large-scale brain networks constructed from functional connectivity (FC) based on temporal synchronization (Van Den Heuvel and Pol 2010, Sporns 2013). This suggests that rBEN shows a more consistent distribution within FC-based brain networks compared to outside of them and constructing rBENN may be meaningful.

In our study, we construct GMVN and rBENN at the individual level according to regional similarity and then evaluate the correlation between rBENN and structural and functional networks from BOLD and MEG. We also assess whether these networks can reflect individual behavioral profiles. The study aims to determine whether individual-based BENN can provide new insights beyond those offered by temporal synchronization brain functional networks and whether they can serve as effective predictors for behavioral profiles.

## 2. Results

### 2.1 The Features and Relationships of Different Brain Networks

We constructed traditional functional connectivity networks (FCN) and resting-state BENN (rsBENN) using resting-state data from the HCP 7T dataset. Additionally, we built movie-watching BENN (mvBENN) using data from 176 participants, with specific methods detailed in the Methods section. Corresponding 3T structural images were used to construct GMVN.

The results show that all networks, including rsBENN, mvBENN, GMVN, and FCN, exhibit higher similarity within the large-scale brain networks (Yeo 7 network) (Yeo, Krienen et al. 2011) than outside of them. Moreover, rsBENN and mvBENN share a similar distribution (Fig. 1A). Fig. 1B shows the average connectivity strength between each node and the global brain across each network. In rsBENN, the average connectivity strength of each node with the whole brain is lower in the default mode network (DMN) and the frontal-parietal network (FPN), while it is higher in the cingulo-opercular network (CON). In mvBENN, the auditory-visual cortex and temporo-parietal junction (TPJ) show lower connectivity, CON exhibits higher connectivity. For GMVN, lower connectivity is observed in the dorsolateral prefrontal cortex (DLPFC) and posterior cingulate cortex/precuneus (PCC/PCu), whereas higher connectivity is found in the dorsomedial prefrontal cortex (DMPFC) and insula. In FCN, the connectivity is lower in the limbic network but higher in the sensorimotor and auditory-visual cortices.

**Fig 1.**
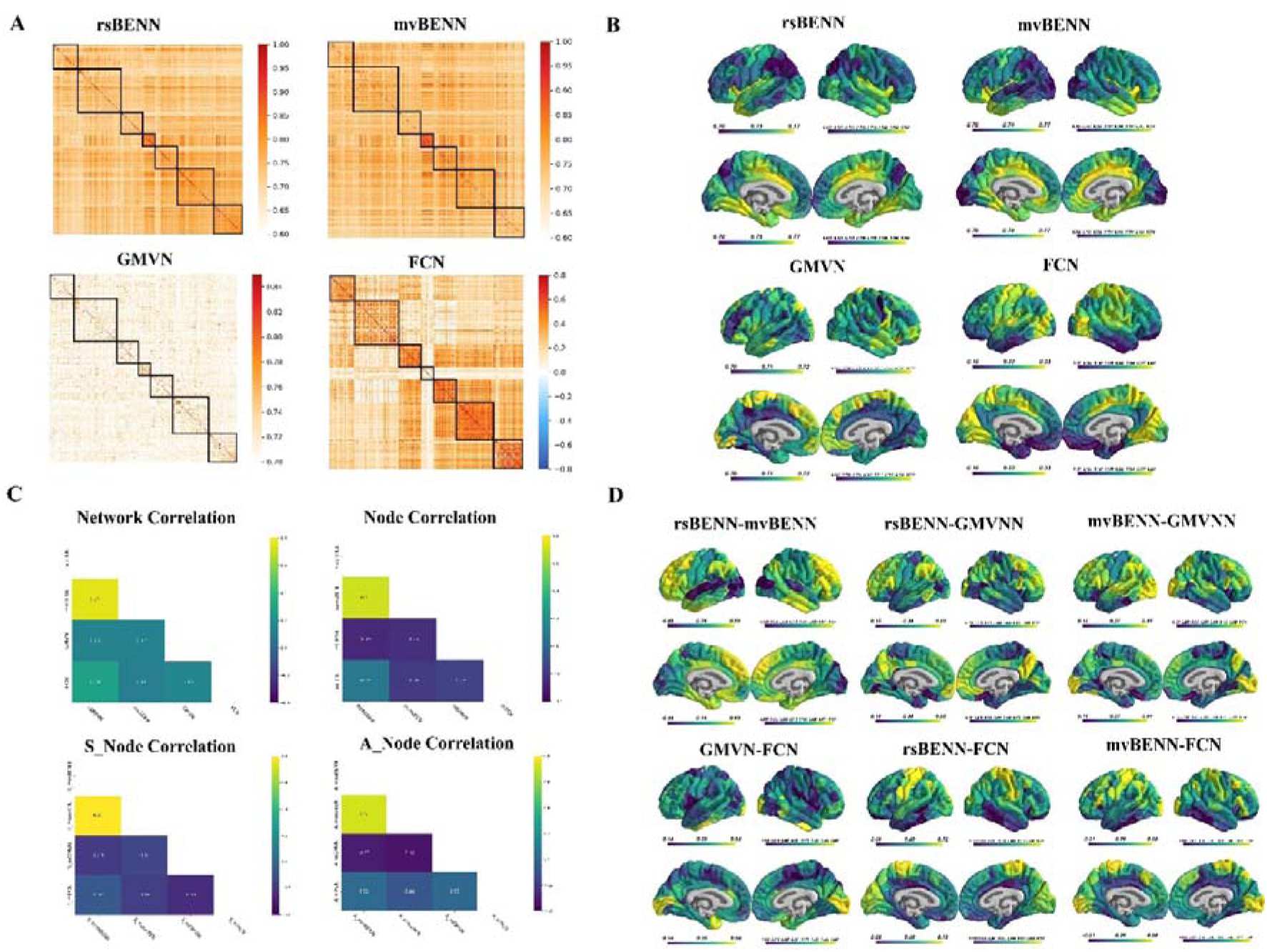
The Features and Relationships of Different Brain Networks. A. The connectivity matrix of the brain network (ordered according to the Yeo–Krienen intrinsic networks: frontoparietal, default mode, dorsal attention, limbic, ventral attention, somatomotor, and visual). B. The average connectivity strength of each node for each brain network. C. The correlations between brain networks. D. The node-wise correlations between brain networks.

We assessed the similarity between networks by performing Pearson correlation analysis on the lower triangular values (Fig. 1C). The correlation between rsBENN and mvBENN was r_(n=79800)_ = 0.74, between rsBENN and GMVN was r_(n=79800)_ = 0.13, between rsBENN and FCN was r_(n=79800)_ = 0.28, between mvBENN and GMVN was r_(n=79800)_ = 0.12, between mvBENN and FCN was r_(n=79800)_ = 0.11, and between GMVN and FCN was r_(n=79800)_ = 0.15. Furthermore, we evaluated the correlation of node connectivity strengths (Fig. 1C). The correlation between rsBENN and mvBENN was r_(n=400)_ = 0.71, between rsBENN and GMVN was r_(n=400)_ = -0.26, between rsBENN and FCN was r_(n=400)_ = 0.08, between mvBENN and GMVN was r_(n=400)_ = -0.23, between mvBENN and FCN was r_(n=400)_ = -0.20, and between GMVN and FCN was r _(n=400)_ = -0.16. The sensorimotor-association (S-A) axis is a crucial feature of the human brain, both in terms of development and functional-structural coupling (Sydnor, Larsen et al. 2021, Rafiei and Rahnev 2022, Liu, Shafiei et al. 2023, Nenning, Xu et al. 2023, Luo, Sydnor et al. 2024, Song 2024). We also analyzed whether the average connectivity strength exhibited different correlations along the S and A axes. We found that the correlation between rsBENN and mvBENN showed no significant difference between the S (r_(n=200)_ = 0.81) and A (r_(n=200)_ = 0.72) regions. However, the correlation between rsBENN and GMVN was more strongly negative in the A region (r = -0.30) than in the S region (r_(n=200)_ = -0.19). No notable difference was observed in the correlation between rsBENN and FCN between the S (r_(n=200)_ = -0.07) and A (r_(n=200)_ = 0.01). The correlation between mvBENN and GMVN was more strongly negative in A (r_(n=200)_ = -0.34) than in the S (r_(n=200)_ = -0.14). The correlation between mvBENN and FCN was more strongly negative in the S (r_(n=200)_ = -0.18) than in the A (r_(n=200)_ = -0.08). Finally, the correlation between GMVN and FCN was more strongly negative in the S (r_(n=200)_ = -0.24) than in the A (r_(n=200)_ = 0.00).

We also evaluated the node-wise correlations between each network (Fig. 1D). The results showed that rsBENN and mvBENN had lower correlations in the sensorimotor cortex, but higher correlations in the association cortex. Similarly, the correlation between rsBENN and GMVN was lower in the sensorimotor cortex and higher in the association cortex. The correlation between mvBENN and GMVN was higher in the visual cortex and FPN, but lower in the insula. The correlation between GMVN and FCN was lower in CON, but higher in the visual cortex. Additionally, rsBENN and FCN showed lower correlations in the association cortex, but higher correlations in the sensorimotor cortex. Similarly, the correlation between mvBENN and FCN was lower in the association cortex and higher in the sensorimotor regions.

### 2.2 Brain network connectome gradient

In recent years, dimensionality reduction of brain networks has become a popular approach for representing the hierarchical structure of brain function using several components, as the connectivity gradient (Margulies, Ghosh et al. 2016, Guell, Schmahmann et al. 2018, Tian, Margulies et al. 2020). In this study, we applied the same approach to analyze the connectivity gradients of each network, extracting the first five gradients for each group of brain networks (Fig 2).

**Fig 2.**
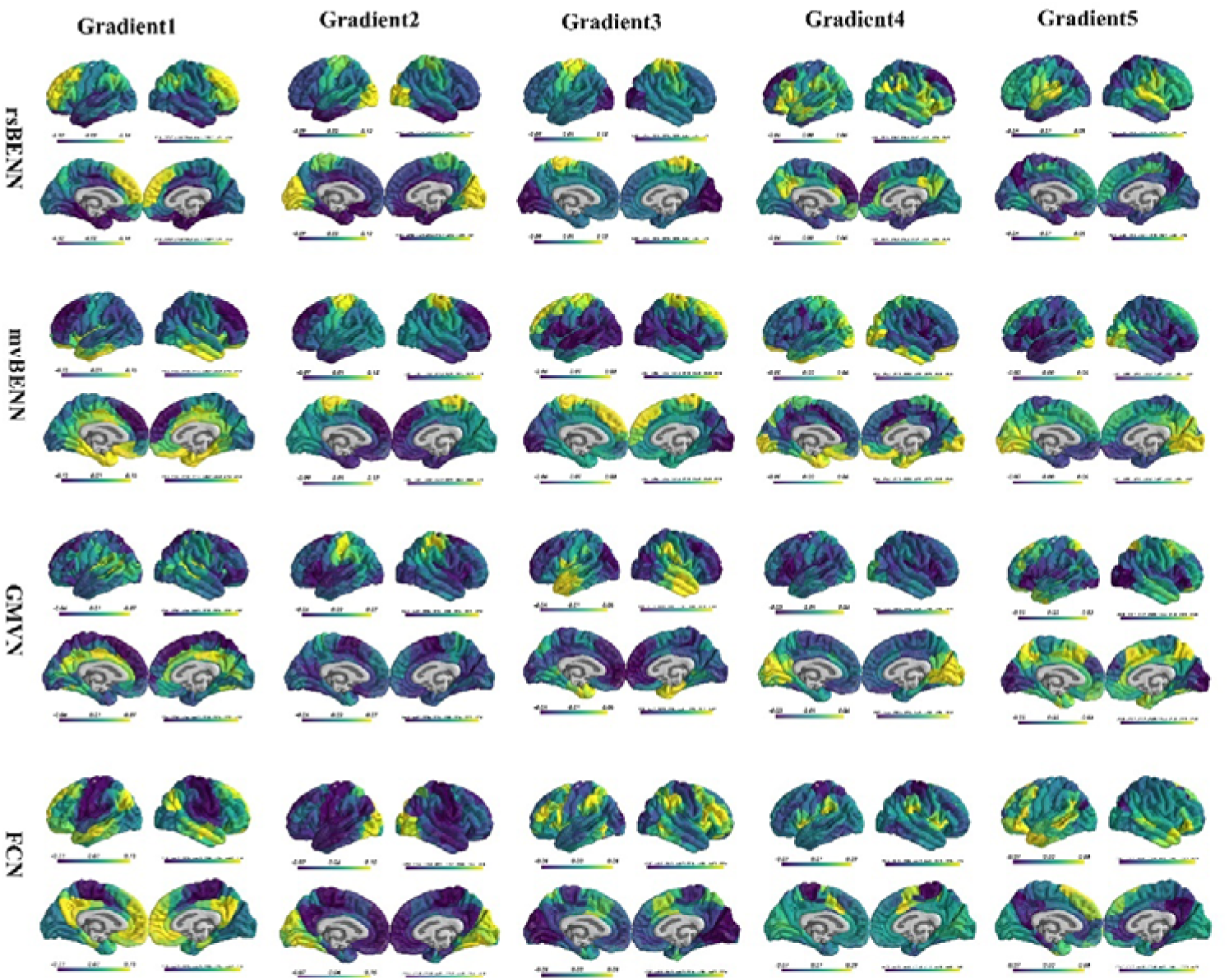
Connectome gradient for each brain network

The results revealed the following connectivity gradient patterns for each network. **For rsBENN:** Gradient 1 showed a pattern of change from the limbic network to the sensorimotor network, and then to the prefrontal cortex. Gradient 2 exhibited a pattern of change from the association cortex to the motor cortex, and then to the visual cortex. Gradient 3 displayed a pattern from the visual cortex to the association cortex, and then to the motor cortex. Gradient 4 revealed a pattern from the prefrontal cortex to the sensorimotor region, followed by the PCC/PCu and insula. Gradient 5 showed a pattern from the default mode network to the visual cortex, and then to the superior temporal gyrus. **For mvBENN:** Gradient 1 presented a pattern from the prefrontal cortex to the sensorimotor region, and then to the limbic network. Gradient 2 exhibited a pattern from the prefrontal cortex to the visual cortex, and then to the motor cortex. Gradient 3 showed a pattern from the visual cortex to the limbic network, and then to the prefrontal cortex. Gradient 4 revealed a pattern from the CON to the prefrontal cortex, and then to the limbic network and visual cortex. Gradient 5 displayed a pattern from the limbic network and temporal lobes to the visual cortex. **For GMVN:** Gradient 1 presented a pattern from the prefrontal cortex to the sensorimotor region, and then to the PCC/PCu. Gradient 2 showed a pattern from the CON to the visual cortex, and then to the sensorimotor region. Gradient 3 revealed a pattern from the prefrontal cortex and visual cortex to the motor cortex, and then to the temporal lobe. Gradient 4 displayed a pattern from the CON to the visual cortex. Gradient 5 exhibited a pattern from the visual cortex and insula to the sensorimotor cortex, followed by the PCC/PCu. **For FCN:** Gradient 1 showed a pattern from the sensorimotor cortex to the visual cortex, and then to the association cortex. Gradient 2 displayed a pattern from the motor cortex to the association cortex, and then to the visual cortex. Gradient 3 presented a pattern from the visual cortex to the sensorimotor cortex, and then to the association cortex. Gradient 4 exhibited a pattern from the motor cortex to the association cortex, and then to the CON. Gradient 5 revealed a pattern from the default mode network to the visual cortex and sensorimotor cortex, and then to the CON.

### 2.3 Connectome-Based Predictive Models

Connectome-based predictive modeling (CPM) is a valuable approach for using brain network information to predict individual behavioral profiles (Shen, Finn et al. 2017). Previous studies based on traditional FCN have demonstrated that CPM can effectively predict traits such as intelligence and emotion (Finn and Bandettini 2021, Wilcox and Barbey 2023). Recently, we have also demonstrated that prediction models based on BEN and GMV show strong predictive ability for cognition and emotion (Song and Wang 2024) . In this study, we constructed CPM based on FCN, GMVN predictive modeling (GMVNPM) based on GMVN, and BENN predictive modeling (BENNPM) based on BENN.

Cognition and emotion scores were obtained from the study (Finn and Bandettini 2021), where principal component analysis (PCA) was performed separately for the cognition and emotion domains. The first principal components were selected as the cognition and emotion scores for their respective domains. We first calculated the correlation between each edge in the network and the cognition and emotion scores. Edges with absolute correlation coefficients greater than 0.2 were selected as predictors. A linear regression model was then used, with 10-fold cross-validation. The correlation between the predicted and actual values was used as the measure of prediction accuracy, and permutation testing with 1,000 repetitions was performed to assess the significance of the correlation coefficients.

The results showed that rsBENN had the highest predictive ability for both cognition and emotion scores (Fig. 3). The prediction accuracies for cognition scores were: rsBENN (r_(n=176)_ = 0.824, *p*<0.001), mvBENN (r_(n=176)_ = 0.609, *p*<0.001), GMVN (r_(n=176)_ = 0.455, *p*<0.001), and FCN (r_(n=176)_ = 0.586, *p*<0.001). For emotion scores, the prediction accuracies were: rsBENN (r_(n=176)_ = 0.593, *p*<0.001), mvBENN (r_(n=176)_ = 0.448, *p*<0.001), GMVN (r_(n=176)_ = 0.555, *p*<0.001), and FCN (r_(n=176)_ = 0.393, *p*<0.001).

**Fig. 3.**
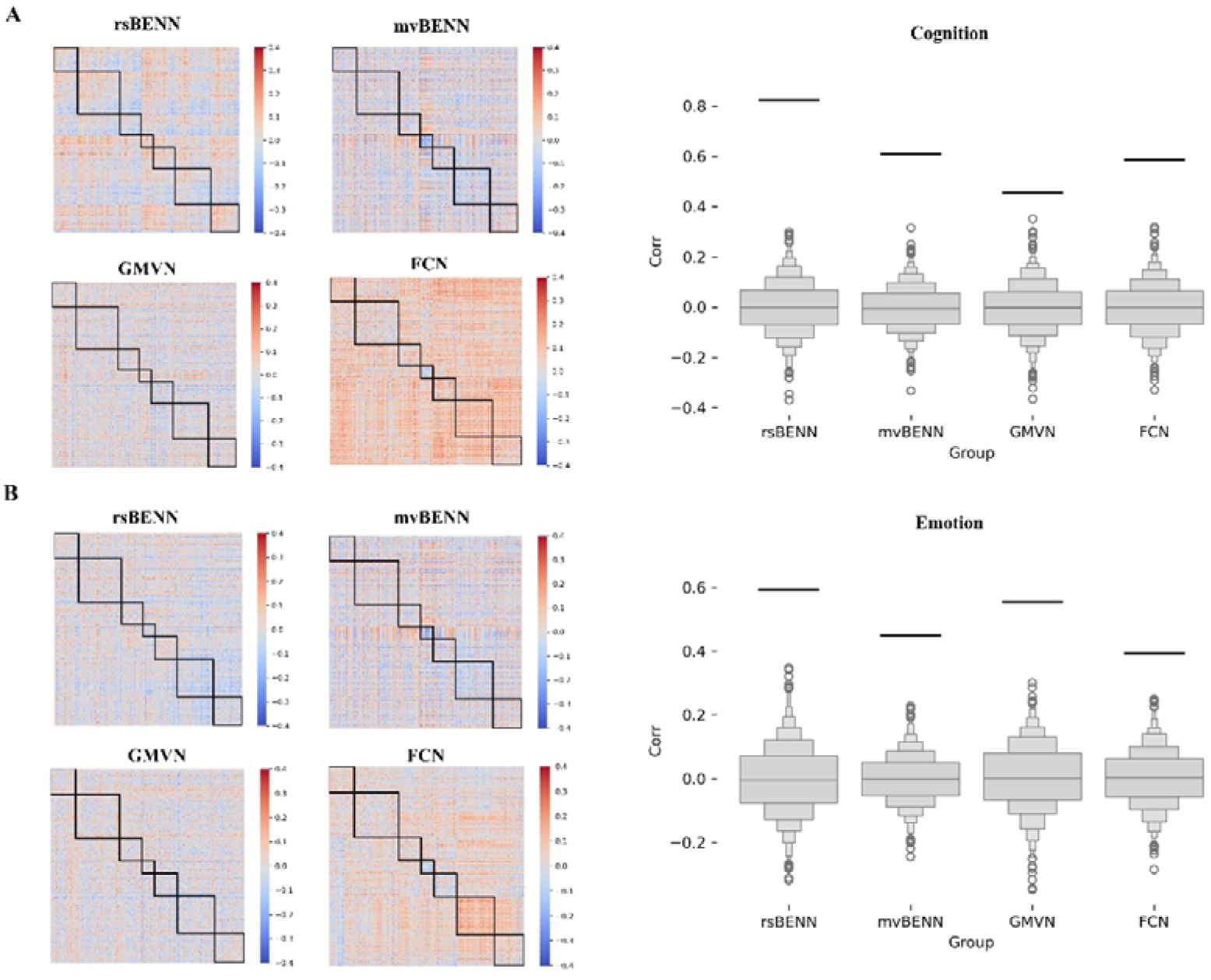
The predictive power of different brain networks for cognition and emotion scores. A. The predictive ability of different networks for cognition scores. B. The predictive ability of different networks for emotion scores. On the left, the Pearson correlation coefficients between the edges of each network and cognition or emotion scores are shown. On the right, the predictive accuracy of each network for cognition or emotion scores is displayed. Accuracy was measured as the Pearson correlation between predicted and observed scores (y-axis). Light gray boxplots represent the null distribution from 10,000 permutations, where cognition or emotion scores and brain networks were randomized across subjects. The black horizontal line indicates the accuracy of the models.

### 3.3 The Relationship with Structural Connectivity Network

We further examined the relationship between BENN and GMVN with the structural connectivity network derived from fiber tracking. The structural connectivity network from (Shafiei, Baillet et al. 2022). We performed correlation analysis on the lower triangular part of the connectivity matrix (Fig. 4B). The results revealed the following correlations: rsBENN with SCN: r _(n=79800)_ = 0.20, mvBENN with SCN: r_(n=79800)_ = 0.17, GMVN with SCN: r_(n=79800)_ = 0.32, FCN with SCN: r_(n=79800)_ = 0.22. Next, we analyzed the correlation between the average connectivity strength of each node. The results were as follows: rsBENN with SCN: r_(n=400)_ = 0.18, mvBENN with SCN: r_(n=400)_ = -0.04, GMVN with SCN: r_(n=400)_ = 0.24, FCN with SCN: r_(n=400)_ = -0.01. Regarding the S-A axis, we found no significant difference in the correlation between rsBENN and SCN across the S-A axis (S: r_(n=200)_ =0.21, A: r_(n=200)_ =0.18). In contrast, the correlation between mvBENN and SCN exhibited an opposite trend along the S-A axis (S: r_(n=200)_ =-0.21, A: r_(n=200)_ =-0.21). The correlation between GMVN and SCN was significant in the S but not in the A (S: r=0.28, A: r_(n=200)_ =0.04). Furthermore, FCN showed a negative correlation with SCN in the S and a positive correlation in the A (S: r_(n=200)_ =-0.14, A: r_(n=200)_ =0.19).

**Fig 4.**
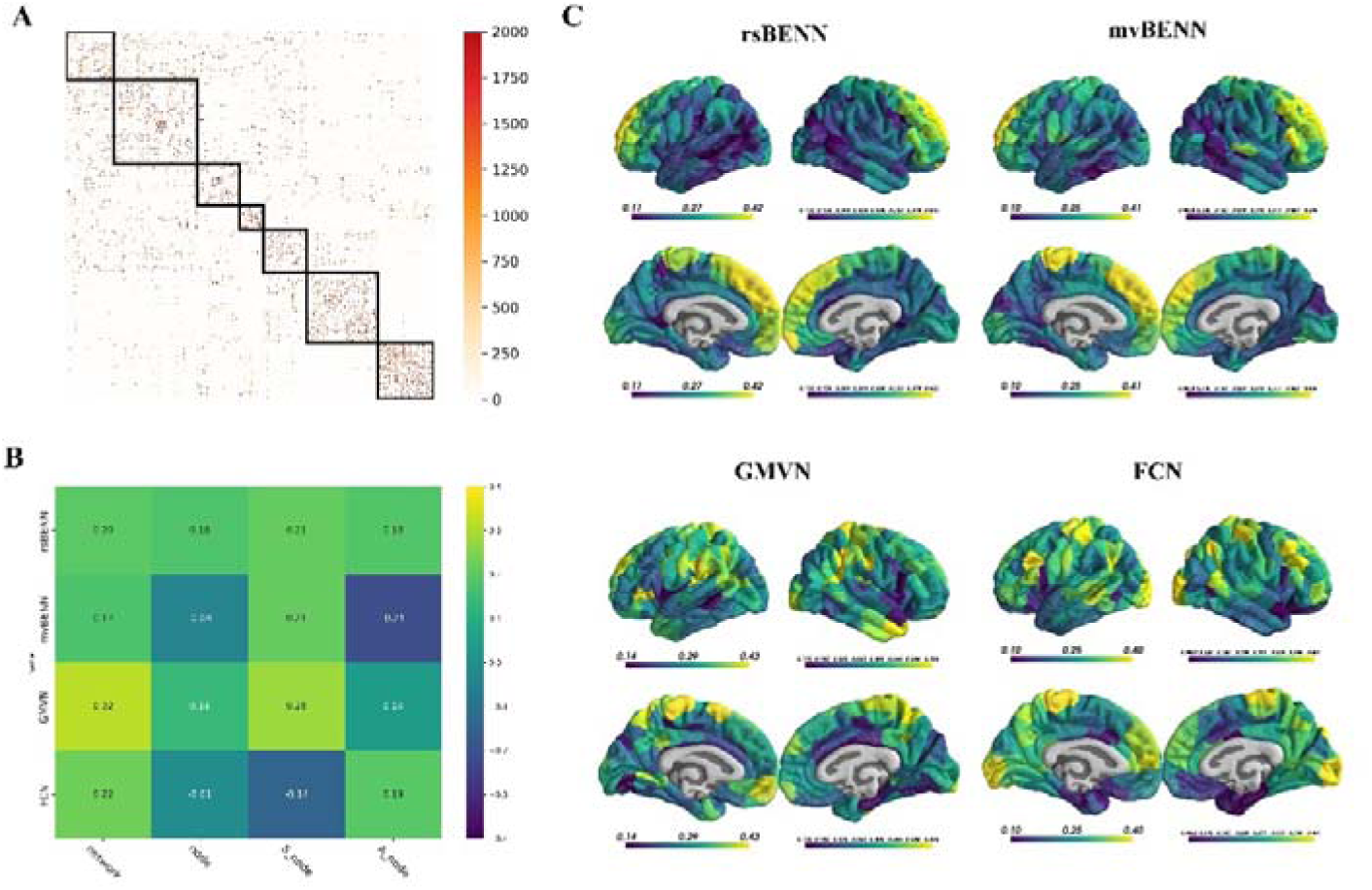
The predictive power of different brain networks for cognition and emotion scores. A. The predictive ability of different networks for cognition scores. B. The predictive ability of different networks for emotion scores. On the left, the Pearson correlation coefficients between the edges of each network and cognition or emotion scores are shown. On the right, the predictive accuracy of each network for cognition or emotion scores is displayed. Accuracy was measured as the Pearson correlation between predicted and observed scores (y-axis). Light gray boxplots represent the null distribution from 10,000 permutations, where cognition or emotion scores and brain networks were randomized across subjects. The black horizontal line indicates the accuracy of the models.

**Fig 5.**
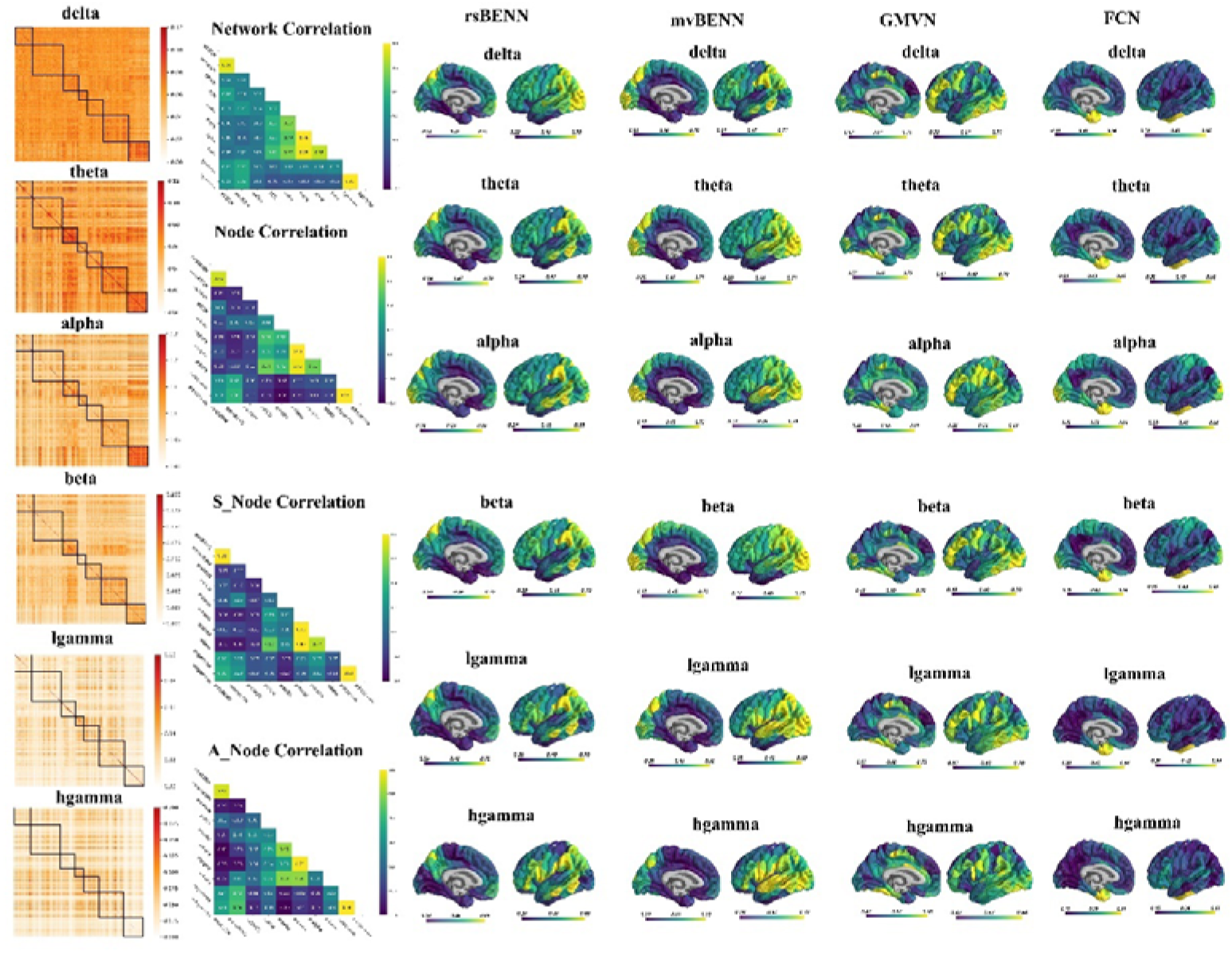
The Relationship with MEG-derived Connectivity Network. From left to right, these are: the MEG-derived Connectivity Network matrix, the correlations between different brain networks, and the node-wise correlations between different brain networks.

We also assessed node-wise correlations across the networks (Fig. 4C). The results indicated that the correlation between rsBENN and SCN was higher in the prefrontal cortex and lower in the temporal lobe. The mvBENN with SCN showed stronger correlations in the prefrontal cortex and weaker correlations in the temporal lobe. GMVN with SCN exhibited higher correlations in the sensorimotor cortex, while the correlation was weaker in the CON. FCN with SCN showed stronger correlations in the sensorimotor and visual cortices, and weaker correlations in the CON and limbic networks.

### 3.4 The Relationship with MEG-derived Connectivity Network

We also assessed the relationship between each network and the MEG-derived connectivity network. The MEG-derived connectivity data were sourced from (Shafiei, Baillet et al. 2022). The MEG-derived networks were divided into six frequency bands: delta, theta, alpha, beta, low gamma, and high gamma. We evaluated the correlations of the lower triangular part of the connectivity matrices for each network. The results showed the following correlation coefficients: rsBEN with delta, theta, alpha, beta, low gamma, and high gamma were 0.17, 0.06, 0.08, 0.05, 0.23, and 0.21, respectively. mvBEN with delta, theta, alpha, beta, low gamma, and high gamma were 0.23, 0.03, -0.01, 0.09, 0.33, and 0.32, respectively. GMVN with delta, theta, alpha, beta, low gamma, and high gamma were 0.14, 0.08, 0.06, 0.08, 0.08, 0.08, and 0.01, respectively. FCN with delta, theta, alpha, beta, low gamma, and high gamma were 0.27, 0.37, 0.32, 0.42, 0.02, and -0.08, respectively.

We also assessed the relationships between the average connectivity strength of each node across networks. The results revealed the following correlations: rsBEN with delta, theta, alpha, beta, low gamma, and high gamma were -0.13, -0.21, -0.10, -0.21, 0.19, and 0.22, respectively. mvBEN with delta, theta, alpha, beta, low gamma, and high gamma were 0.05, -0.28, -0.27, -0.13, 0.32, and 0.38, respectively. GMVN with delta, theta, alpha, beta, low gamma, and high gamma were 0.02, -0.16, -0.16, -0.13, -0.09, -0.13, respectively. FCN with delta, theta, alpha, beta, low gamma, and high gamma were 0.18, 0.44, 0.35, 0.54, -0.20, and -0.25, respectively. When analyzing according to the S-A axis, we observed the following correlations in the S region: rsBEN with delta, theta, alpha, beta, low gamma, and high gamma were -0.06, -0.26, -0.11, -0.31, 0.28, and 0.35, respectively. mvBEN with delta, theta, alpha, beta, low gamma, and high gamma were 0.15, -0.28, -0.15, -0.32, 0.16, and 0.23, respectively. GMVN with delta, theta, alpha, beta, low gamma, and high gamma were -0.17, -0.23, -0.21, -0.23, -0.08, and -0.09, respectively. FCN with delta, theta, alpha, beta, low gamma, and high gamma were 0.02, 0.26, 0.15, 0.52, -0.05, and -0.06, respectively. In the A region, the correlations were as follows: rsBEN with delta, theta, alpha, beta, low gamma, and high gamma were -0.26, -0.42, -0.36, -0.27, 0.24, and 0.30, respectively. mvBEN with delta, theta, alpha, beta, low gamma, and high gamma were 0.04, -0.20, -0.35, 0.16, 0.42, and 0.46, respectively. GMVN with delta, theta, alpha, beta, low gamma, and high gamma were 0.18, -0.02, -0.04, 0.02, -0.18, and -0.27, respectively. FCN with delta, theta, alpha, beta, low gamma, and high gamma were 0.16, 0.28, 0.14, 0.32, 0.02, and -0.01, respectively.

The node-wise correlation analysis between networks revealed the following results: rsBENN showed higher correlations with the MEG-derived connectivity network in the TPJ, and lower correlations in the limbic network and CON. The mvBENN had stronger correlations with the MEG-derived connectivity network in the visual cortex and temporoparietal junction, and weaker correlations in the limbic network and CON. GMVN exhibited higher correlations with the MEG-derived connectivity network in the dorsolateral prefrontal cortex, and lower correlations in the medial prefrontal cortex. FCN showed higher correlations with the MEG-derived connectivity network in the temporal pole, and lower correlations in the CON, DMN, and FPN.

## 3. Discussion

Our study demonstrates that brain networks constructed using simple regional similarity, both at the structural and functional brain network levels, reveal potential physiological significance. First, these constructed networks are partially correlated with traditional structural and functional networks, yet their features cannot be fully captured by conventional networks, suggesting they provide additional information beyond what traditional networks offer. Second, gradient analysis also revealed meaningful patterns. Third, when it comes to predicting behavioral phenotypes, the rsBENN showed the highest predictive power, indicating its potential superiority in behavioral and clinical disease prediction.

The morphological brain networks were constructed by Kullback-Leibler (KL) divergence have already demonstrated their significance for potential application to individual differences studies (Kong, Wang et al. 2014, Wang, Jin et al. 2016). In this study, we employed a simpler and more computationally efficient method. Moreover, we extended this approach to functional brain networks, revealing information that traditional FCN are unable to capture, and even achieving higher predictive accuracy for behavioral traits. Using similar methods to construct individual brain networks could provide insights from structural indicators such as myelination, cortical thickness, and other functional metrics like amplitude of low-frequency fluctuation (Zang, Jiang et al. 2004, Yu-Feng, Yong et al. 2007, Zou, Zhu et al. 2008), regional homogeneity (Zang, Jiang et al. 2004), and task-related activations.

In summary, this study systematically evaluates the construction of brain network features from regional brain profiles and their potential applications. It lays the groundwork for understanding brain function from local to network levels and paves the way for future applications.

## 4. Methods

### 4.1 Dataset

The BEN maps were calculated from preprocessed resting-state fMRI and movie-watching fMRI from HCP 7T release (Van Essen, Smith et al. 2013). The BEN maps used in this study are the same as that in our previous research (Song and Wang 2024, Song and Wang 2024, Song and Wang 2024), with a total of 176 participants included. For each participant, the resting-state BEN (rsBEN) and movie-watching BEN (mvBEN) were segmented into 400 parcels based on the Schaefer 400 (Schaefer, Kong et al. 2018). The group-averaged structural networks and functional brain networks based on MEG were obtained from https://github.com/netneurolab/shafiei_megfmrimapping/tree/main/data that were derived from 33 participants in the HCP Young Adult. For preprocessing and network construction of this data, please refer to the original paper (Shafiei, Baillet et al. 2022). Additionally, we also computed functional brain networks using zero-lag functional connectivity based on the 176 participants using the 400 regions based on HCP 7T REST2 AP data.

### 4.2 Brain network construction

#### 4.2.1 BENN construction

The 400 regions as nodes and the absolute difference between the z-score of the average BEN of each region as edge, resulting in a 400×400 matrix for each participant. Then, the matrix was scaled into [0,1], finally subtracting each normalized element from 1, which is then used to represent the connection strength between regions.

#### 4.2.2 GMVN construction

The construction method is the same as that of BENN, with the difference being that the BEN values are replaced with GMV values.

#### 4.2.3 FCN construction

The Pearson correlation was calculated for the time series across 400 nodes, and the correlation coefficients were then Fisher z transformed. The z-score intensity represents the connection strength of the edges.

## Acknowledgments

MRI and MEG data were provided by the Human Connectome Project, WU-Minn Consortium (Principal Investigators: David Van Essen and Kamil Ugurbil; 1U54MH091657) funded by the 16 NIH Institutes and Centers that support the NIH Blueprint for Neuroscience Research; and by the McDonnell Center for Systems Neuroscience at Washington University. We thank Golia Shafiei et al for sharing their dataset.

## Notes

In this manuscript, I systematically evaluate the comparison of features between individual brain networks constructed using BEN and GMV, as well as their predictive capabilities for behavior phenotypes. The results highlight the strong potential of BENN and GMVN in bridging the understanding of brain function from the local to the network level. This sets the foundation for future research on brain development, individual differences, and clinical applications. However, due to current computational limitations and time constraints, several aspects remain unfinished. Upon completing my doctoral thesis this year, which is a distinct project from BEN, I will further refine and expand upon this series of BEN studies.

